# Jasmonic acid signaling and glutathione coordinate plant recovery from high light stress

**DOI:** 10.1101/2024.10.16.618759

**Authors:** Mehmet Kılıç, Peter J. Gollan, Eva-Mari Aro, Eevi Rintamäki

**Affiliations:** Plant Molecular Biology, Department of Life technologies, University of Turku, 20014 Turku, Finland

## Abstract

High light-induced chloroplast retrograde signaling originates from the photosynthetic apparatus and regulates nuclear gene expression to enhance photoprotection and coordination of cell metabolism. Here, we analyzed the transcript profiles and accumulation of ROS, stress hormones and small molecule antioxidants to investigate the signaling mechanisms operating under HL stress, and in particular during plant recovery under growth conditions. Exposure to high light for 15 min induced a number of ^1^O_2_- and H_2_O_2_-responsive genes and accumulation of an oxidative form of glutathione and ascorbate, the hallmarks of oxidative stress in cells. Prolonged exposure to high light resulted in accumulation of transcripts encoding oxylipin biosynthesis enzymes, leading to accumulation of 12-oxo-phytodienoic acid and jasmonic acid. However, the expression of several jasmonic acid -responsive genes, already induced by HL, peaked during the recovery together with accumulation of jasmonic acid, reduced glutathione and ascorbate, highlighting the critical role of jasmonic acid signaling in restoring chloroplast redox balance following high light stress. The involvement of jasmonic acid signaling in recovery-sustained gene expression was further confirmed by conducting experiments with jasmonic acid receptor mutants. High light exposure of only two min was sufficient to induce some recovery-sustained genes, indicating rapid response of plants to changing light conditions. We propose that ROS production at high light induces the signaling cascade for early oxylipin biosynthesis and 12-oxo-phytodienoic acid accumulation, while increased accumulation of jasmonic acid in recovery-phase activates the genes that fully restore the glutathione metabolism, and ultimately allow recovery from short-term high light stress.

## Introduction

Photosynthesis plays a crucial role in sensing environmental cues and relaying signals for regulation of plant acclimation (Walters, 2005; Gollan et al., 2015; Morales and Kaiser, 2020). Photosynthetic machinery converts environmental cues into biochemical messengers that adjust the expression of genes in both the cell nucleus and plastids. This acclimation response helps the organism cope with stresses and to acclimate to alternating environments. Photosynthetic light reactions, occurring in photosystem (PS) II and PSI together with their light-harvesting complexes (LHC), collect and transform light energy to chemical energy to fuel the assimilation of atmospheric CO_2_ into organic energy-rich compounds. Environmental stress, such as high light (HL), can disrupt the electron flow, leading to production of reactive oxygen species (ROS) and reactive electrophile species (RES) (Farmer and Davoine, 2007; Khorobrykh et al., 2020; Fitzpatrick et al., 2022) which, if not controlled, are hazardous to biological molecules. In addition to cellular damage, ROS and RES also initiate signals to protect and acclimate the photosynthetic machinery from deleterious effects of redox changes in the chloroplast.

Oxylipins are important redox components regulating growth and stress responses in plants (recent review, see Knieper et al., 2023). ROS induces the synthesis of oxylipins, including 12-oxo-phytodienoic acid (OPDA) and jasmonic acid (JA). They are derivatives of oxygenated α-linolenic acid released from plastid membranes by lipases (DAD1, DALL1), and are synthesized in an enzymatic pathway (Farmer and Davoine, 2007; Wasternack and Song, 2017; Knieper et al., 2023). OPDA synthesis starts with the oxygenation of α-linolenic acid by 13-LIPO-OXYGENASE (LOX) producing 13-hydroperoxy-octadecatrienoic acid (13-HPOT), which is then processed to OPDA by ALLENE OXIDASE SYNTHASE (AOS) and ALLENE OXIDASE CYCLASE (AOC) (Schaller and Stintzi, 2009; Wasternack and Song, 2017). OPDA is transported from plastids to peroxisomes to synthesize JA by 12-OXOPHYTODIENOATE REDUCTASE 3 (OPR3) followed by three cycles of β-oxidation catalyzed by ACYL-COA OXIDASE (ACX1) (Schaller and Stintzi, 2009; Wasternack and Song, 2017). *JAR1* gene is required for biological activation of JA via conjugation with isoleucine (JA-Ile) (Suza and Staswick, 2008; Wasternack and Song, 2017).

Both OPDA and JA function as signaling molecules modifying gene expression in the nucleus (Chini et al., 2007; Wasternack and Song, 2017). Oxylipins that contain α,β-unsaturated carbonyl bonds (e.g. OPDA) are categorized as RES due to their inherent reactivity with thiol groups of cellular proteins (Améras et al., 2003; Farmer and Davoine, 2007). OPDA and JA were observed to take part in antioxidant defense response by inducing antioxidant gene expression (Sasaki-Sekimoto et al., 2005; Gollan and Aro, 2020). JA is sensed by its receptor CORONATINE INSENSITIVE 1 (COI1) (Wasternack and Song, 2017) that is a subunit of the E3 ubiquitin protein ligase (SKP1-Cullin-F-box, SCF^COI1^) (Yan et al., 2013; Wasternack and Song, 2017). SCF^COI1^ labels proteins with ubiquitin to induce the proteolytic degradation (Chini et al., 2007; Yan et al., 2013). JA-Ile binds to the SCF^COI1^ complex and facilitates the ubiquitination and degradation of the JAZ transcriptional repressors, allowing transcription factors to bind to JA-responsive genes to initiate transcription (Chini et al., 2007; Dombrecht et al., 2007). Via this induction mechanism, JA signaling is involved in regulation of CO_2_ fixation, antioxidant metabolism, degradation of chlorophylls, reallocation of resources from growth toward defense systems, and senescence (Sasaki-Sekimoto et al., 2005; Shan et al., 2011; Park et al., 2013; Zhang et al., 2020).

Cells harbor both enzymatic and non-enzymatic ROS scavenging systems to protect against excessive damage by ROS. Enzymes, such as SUPEROXIDE DISMUTASEs (SOD), CATALASEs (CAT), ASCORBATE PEROXIDASEs (APX), and GLUTATHIONE PEROXIDASEs (GPX) directly detoxify ROS, whereas other enzymes such as THIOREDOXINs (TRX), GLUTAREDOXINs (GRX), and PEROXIREDOXINs (PRX) re-reduce proteins oxidized by ROS. Non-enzymatic ROS scavenging system includes molecular antioxidants such as ascorbate, tocopherols, and glutathione, which act as electron donors to neutralize ROS. Glutathione exhibits a high affinity to both ROS and RES (Dixon and Edwards, 2009; Foyer and Noctor, 2011; Ito and Ohkama-Ohtsu, 2023), thereby preserving the cellular integrity. Glutathione exists in two forms: the reduced (GSH) and oxidized disulfide (GSSG). In the presence of oxidants, electrons from GSH are utilized to neutralize ROS with concomitant generation of GSSG. Consequently, under oxidative stress, the ratio of GSH to GSSG significantly decreases (Toldi et al., 2019; Gasperl et al., 2021) but after relaxation of stress, oxidized GSSG is reduced back to GSH by GLUTATHIONE REDUCTASE (GR) and NADPH.

Induction of protective gene expression and antioxidant metabolism have been extensively studied in HL-treated plants (Hernández et al., 2004; Chan et al., 2016; Balfagón et al., 2019; Huang et al., 2019; Alvarez-Fernandez et al., 2021), while far less is known about their fate after transferring the plants back to growth light (GL) conditions for recovery (R) (Crisp et al., 2017). Furthermore, short-term HL treatments ranging from minutes up to an hour, do not yet induce acclimation processes but modify cell metabolism, helping the plant to cope under changing light conditions (Gollan et al., 2023). In this study, we analyzed global gene expression changes, together with the stress hormone and antioxidant levels, in leaves exposed to short-term HL stress and during subsequent R from the stress at GL. We show that JA is an important mediator of nuclear gene expression in leaves during R from HL stress and that glutathione helps to balance the redox state of the cell when the HL illumination is terminated.

## Results

### JA, OPDA and GSH are involved in regulation of gene expression during recovery from HL treatment

Global gene expression analysis was performed after the HL treatments of 15 and 60 min (HL15 and HL60) and after subsequent R phases in GL for 15 (R15) and 60 min (R60), and the results were expressed with respect to GL control without any HL treatment. Distinct patterns of differential gene expression were identified in each phase. A total of 6998 differentially expressed genes (DEGs), compared to the GL control, were detected in the experiment (Figure S1). The number of DEGs was substantially higher during the R compared to the preceding HL treatment. Specifically, the highest number of DEGs was observed at R60, while the lowest was recorded at HL15 (Figure S1A). This indicated that considerable reprogramming and adjustment occur in the cell during R phases after HL treatment.

We used the DEGs to identify, at the transcript level, the various biological processes that were affected during the HL and R phases of the experiment. Gene Ontology (GO) terms related to ROS responses were particularly enriched in genes upregulated during the HL treatment (Table S1). Transcripts of glutathione metabolism and JA signaling, on the other hand, accumulated during HL60 as well as R15 and R60 (Table S1). GO terms related to abscisic acid (ABA) signaling were moderately upregulated in HL15, whereas GO terms related to salicylic acid (SA) signaling were upregulated in both HL and R, especially in HL15 and R15 (Table S1).

Since transcripts of hormone and oxidative stress-related biological processes were found to be enriched GO analysis (Table S1), we next analyzed our gene expression data with respect to known responses induced by metabolites such as ^1^O_2_, H_2_O_2_, ABA, OPDA, JA and SA (Op Den Camp et al., 2003; Xin et al., 2005; Gollan and Aro, 2020; Zhang et al., 2020). Most of OPDA and JA biosynthesis genes were slightly upregulated already in HL15, but their expression was further increased in HL60 and remained high or increased in R phase (Figure 1A). A similar trend was also observed for other JA- and OPDA-responsive genes (Figures 1B and 1C). The expression of ^1^O_2_-responsive genes peaked in R15 (Figure S2A), whereas the expression of H_2_O_2_-responsive genes increased with HL treatment and decreased during the R phase (Figure S2B). About half of the ABA-responsive genes showed increased expression in HL60 and R15, while the expression of SA-responsive genes did not significantly change in HL or during R (Figures S3A and S3B).

**Figure 1:**
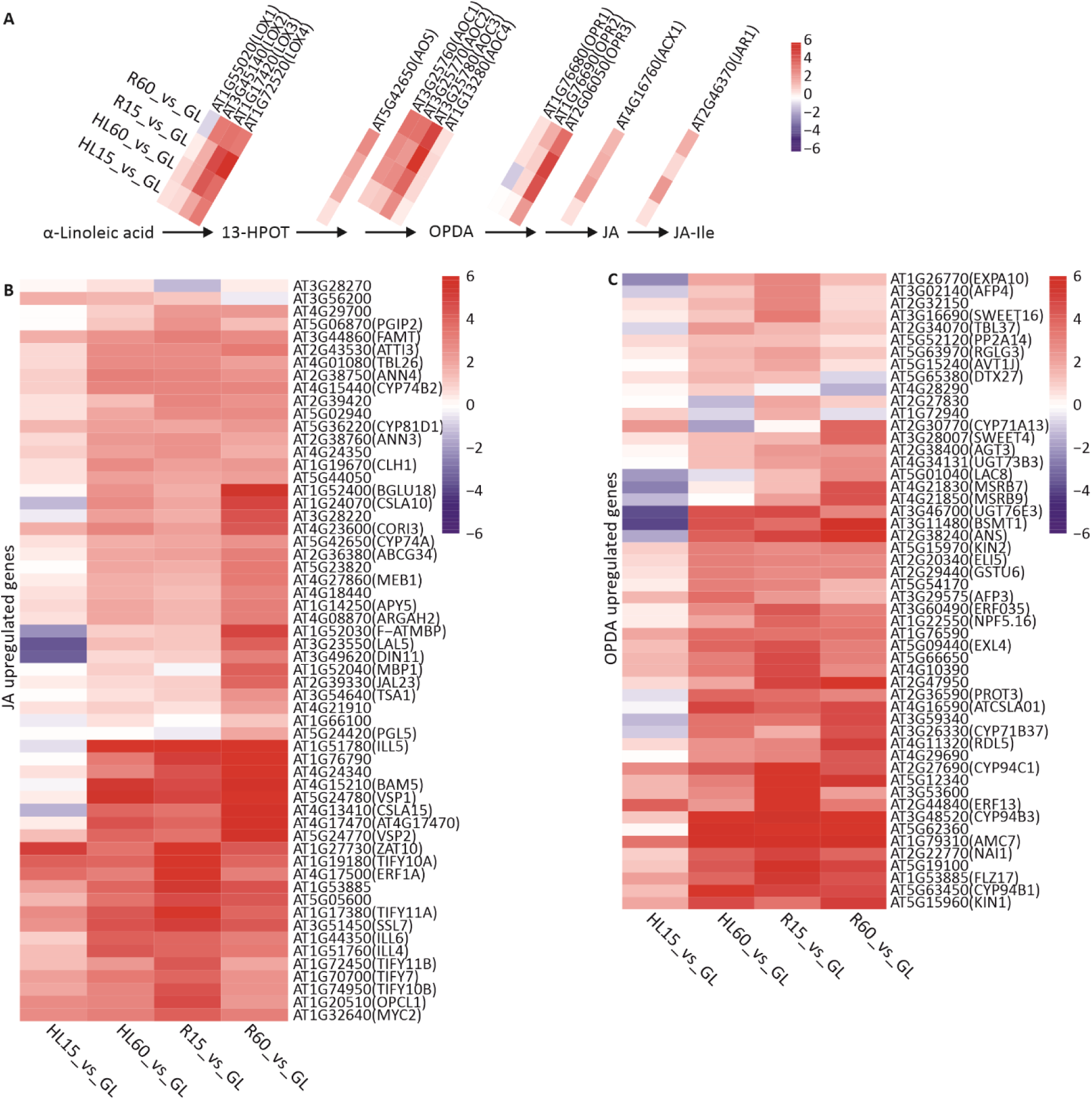
Differential expression of oxylipin responsive genes in leaves treated in high light (HL) and in recovery (R) at growth light (GL) in comparison to control GL leaves. HL treatment was performed by exposing plants to HL for 15 (HL15) and 60 min (HL60), while recovery was performed by transferring HL60 samples to GL to recover for 15 (R15) and 60 (R60) min. A) Differential expression of 12-oxo-phytodienoic acid (OPDA) and jasmonic acid (JA) synthesis genes. For the gene names, see the text. B) Differential expression of JA-responsive genes. C) Differential expression of OPDA-responsive genes. The genes indicated in the figure B and C are reported to be upregulated in response to OPDA and JA treatments in (Gollan and Aro, 2020). Red and blue color scale (log2 fold change) shows the degree of upregulation and downregulation of the genes, respectively.

We also analyzed, after both the HL and R phases, the expression response of genes encoding proteins involved in GSH metabolism. The genes related to GSH biosynthesis, cellular transportation and degradation were differentially expressed in HL and subsequent R phases (Figure 2, Table S2). The expression levels of *GLUTATHIONE SYNTATHASE 2 (GSH2)* and *GAMMA-GLUTAMYL CYCLOTRANSFERASE 2 (GGCT2s)* doubled during R60 in comparison to GL (Figure 2). *GSH2* encodes the biosynthetic enzyme that catalyzes the addition of glycine (Gly) to y-Glu-Cys, while *GGCT2s* encodes the enzymes breaking down and recycling GSH (Dorion et al., 2021; Ito and Ohkama-Ohtsu, 2023). Conversely, the expression of the *OLIGOPEPTIDE TRANSPORTER (OPT)* and *GAMMA-GLUTAMYL TRANSPEPTIDASE 1 (GGT1)* genes, encoding proteins involved in transporting GSH to the apoplast and breaking down GSSG in the apoplast (Zhang et al., 2016; Wongkaew et al., 2018; Ito and Ohkama-Ohtsu, 2023), respectively, was halved in both the HL and R treatments in comparison to GL (Figure 2). Furthermore, the expression of the genes related to GSH metabolism was enhanced both in the HL and R phases (Figure 2). GSH is exported from chloroplasts to cytosol via CHLOROQUINE-RESISTANCE TRANSPORTER (CRT)-LIKE TRANSPORTER (CLT) and in the cytosol, the GLUTATHIONE S-TRANSFERASE (GSTU) catalyzes conjugation of GSH with OPDA (GS- OPDA; Skipsey et al., 2011). Seven out of 11 *GSTU* genes were upregulated in response to HL and R treatments (Figure 2). Additionally, the *GAMMA-GLUTAMYL TRANSPEPTIDASE 4* (*GGT4*) gene was highly upregulated by HL and R treatments (Figure 2), indicating that the GS-OPDA conjugate produced in cytosol, is transported to the vacuole via MULTIDRUG RESISTANCE-ASSOCIATED PROTEIN (MRP) transporters for processing by GGT4 (Dorion et al., 2021; Ito and Ohkama-Ohtsu, 2023). Thus, the DEGs related to GSH metabolism suggest that the biosynthesis and recycling of GSH are critical for both HL and R processes (Figure 2).

**Figure 2:**
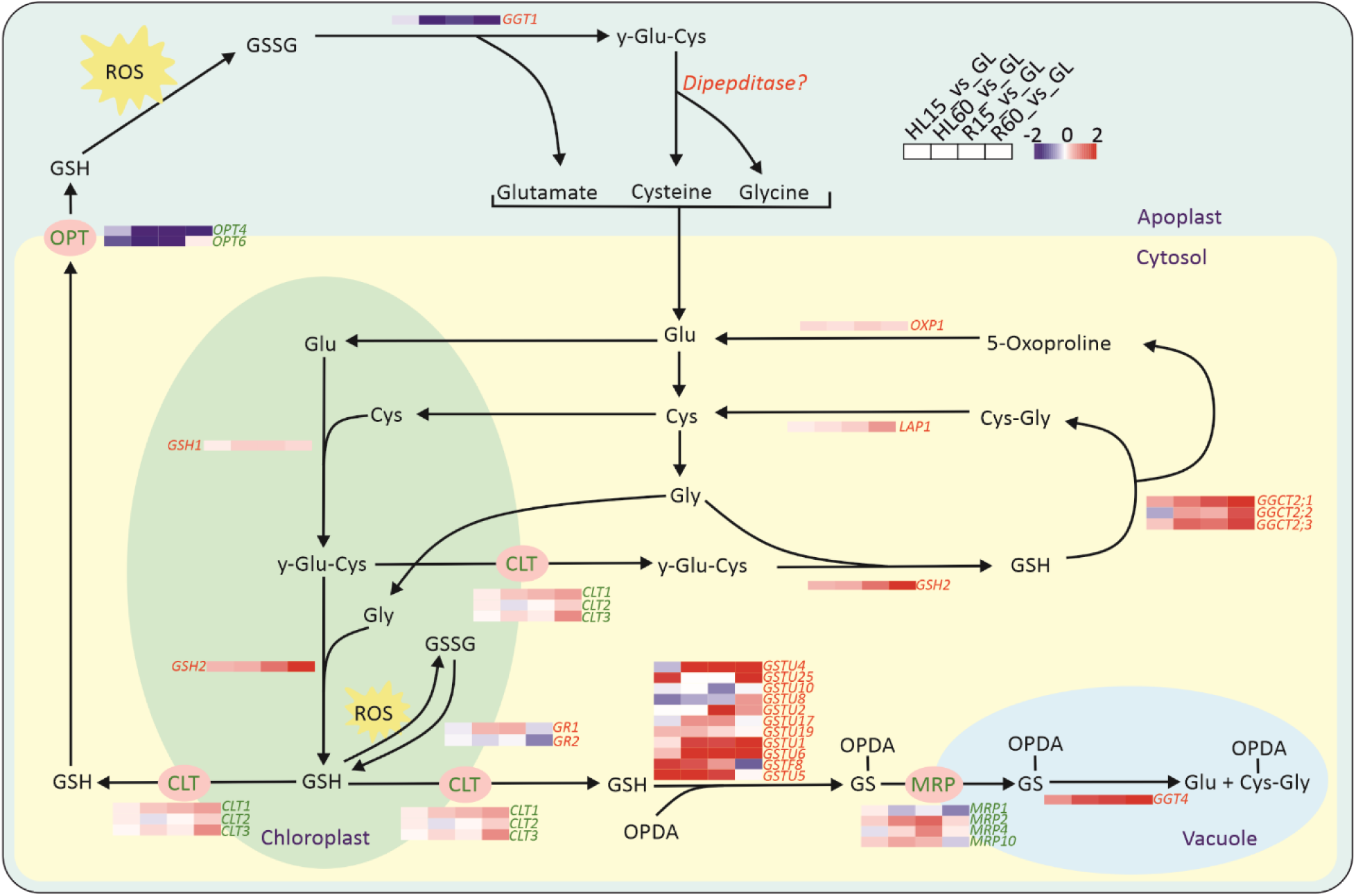
Differential expression of genes involved in glutathione metabolism in leaves treated in high light (HL) and recovery (R) at growth light (GL) in comparison to control GL leaves. HL treatment was performed by exposing plants to HL for 15 (HL15) and 60 min (HL60), while recovery was performed by transferring HL60 samples to GL to recover for 15 (R15) and 60 min (R60). The figure of GSH metabolism was modified from (Dorion et al., 2021). Red and blue color scale (log2 fold change) shows the degree of upregulation and downregulation of the genes, respectively. For the gene names, see the text. For the accession numbers of the genes and fold changes, see Table S2.

JA-responsive genes were activated at HL60, and they remained upregulated, or their transcripts further increased during the R phase (Figure 1). These genes are called R-sustained genes hereafter. Since H_2_O_2_ treatment has been shown to trigger JA biosynthesis in plant leaves (Hieno et al., 2019; Lv et al., 2019), we next investigated whether the representative R-sustained genes are induced by H_2_O_2_ treatment alone at GL condition similar to that in HL exposure. To this end, the genes *VSP2*, *JAZ8*, *Rap2.6*, *AOC2* and *AOS,* which are highly induced during the R phase (Table S3), were selected as marker genes for qPCR, and the upregulated expression of *HSP* genes (Figure 3A), which is known to be induced by H_2_O_2_ (Gollan & Aro, 2020), ensured the successful penetration of H_2_O_2_ into the cells. The response to external H_2_O_2_ of the selected R-sustained genes varied but mainly followed the expression pattern observed in HL- and subsequent R-treated leaves. *JAZ8* gene was induced during the H_2_O_2_ treatment, and its expression continued after removal of H_2_O_2_, whereas *AOC2*, *Rap2.6* and *AOS* genes were upregulated afterwards in GL during recovery from the H_2_O_2_ treatment (Figure 3B). The experiment suggests that these genes respond to JA synthesized in the H_2_O_2_-treated leaves. *VSP2* gene did not respond to this treatment at all.

**Figure 3:**
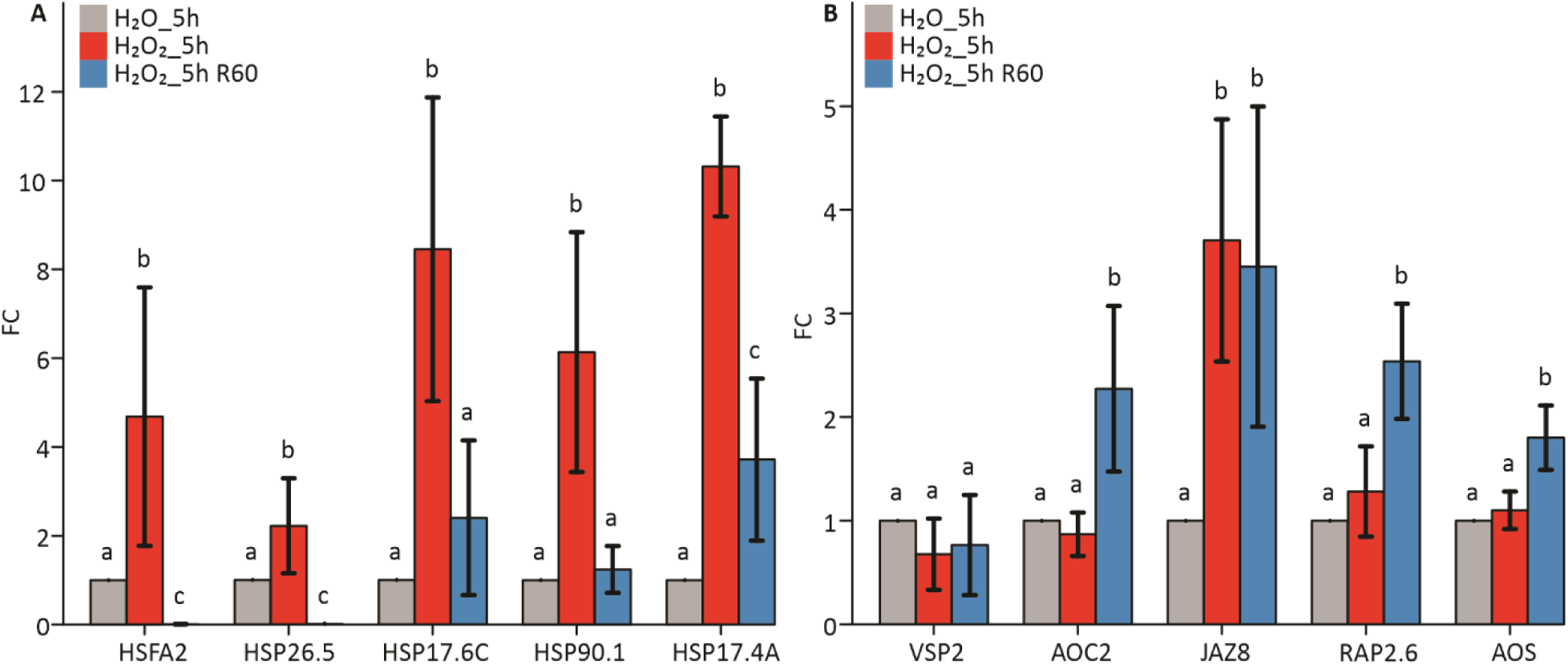
Relative expression of A) *HSP* genes and B) recovery-sustained genes in response to the H_2_O_2_ treatment and recovery in water after the H_2_O_2_ treatment. qPCR test was performed with leaves that were floated on water for 5h (H_2_O_5h), or on 20 mM H_2_O_2_ for 5h (H_2_O_2__5h) at growth light (GL). After the treatment, the leaves were transferred to water for recovery for 60 min (H_2_O_2__5h R60). The following (A) HSP genes (see Table S3): *HSFA32*, *HSP26.5*, *HSP17.6C*, *HSP90.1* and *HSP17.4A*, and (B) R-sustained genes: *VSP2*, *AOC2*, *JAZ8*, *Rap2.6* and *AOS,* were analyzed. The fold change (FC) of gene expression was normalized to control values (H_2_O_5h). Values represent the mean ± SD of four samples. The letters indicate significant differences between the treatments (ANOVA, Tukey-HSD, p < 0.05).

To confirm that JA is indeed involved in induction of R-sustained genes, we further tested the expression of a specific set of R-sustained marker genes in mutants lacking the JA receptor COI1 (*coi1-1*, *coi1-2*) using qPCR analysis. In this experiment, the leaves of wild type (WT) and the two *coi1* mutants were collected from GL and after the exposure of leaves first to HL treatment for 60 min and then for 60 min to GL for recovery (R60; Figure 4). Under initial GL conditions, the expression of selected genes was either strongly (*VSP2*, *JAZ10* and *Rap2.6*) or slightly (AOS) lower in both *coi1* mutants compared to WT (Figure 4). After HL treatment of 60 min followed by subsequent R for 60 min, the marker genes were significantly upregulated in WT, but not in the *coi1* mutants (Figure 4), providing strong evidence that the JA signaling indeed initiates the expression of these genes during the R phase.

**Figure 4:**
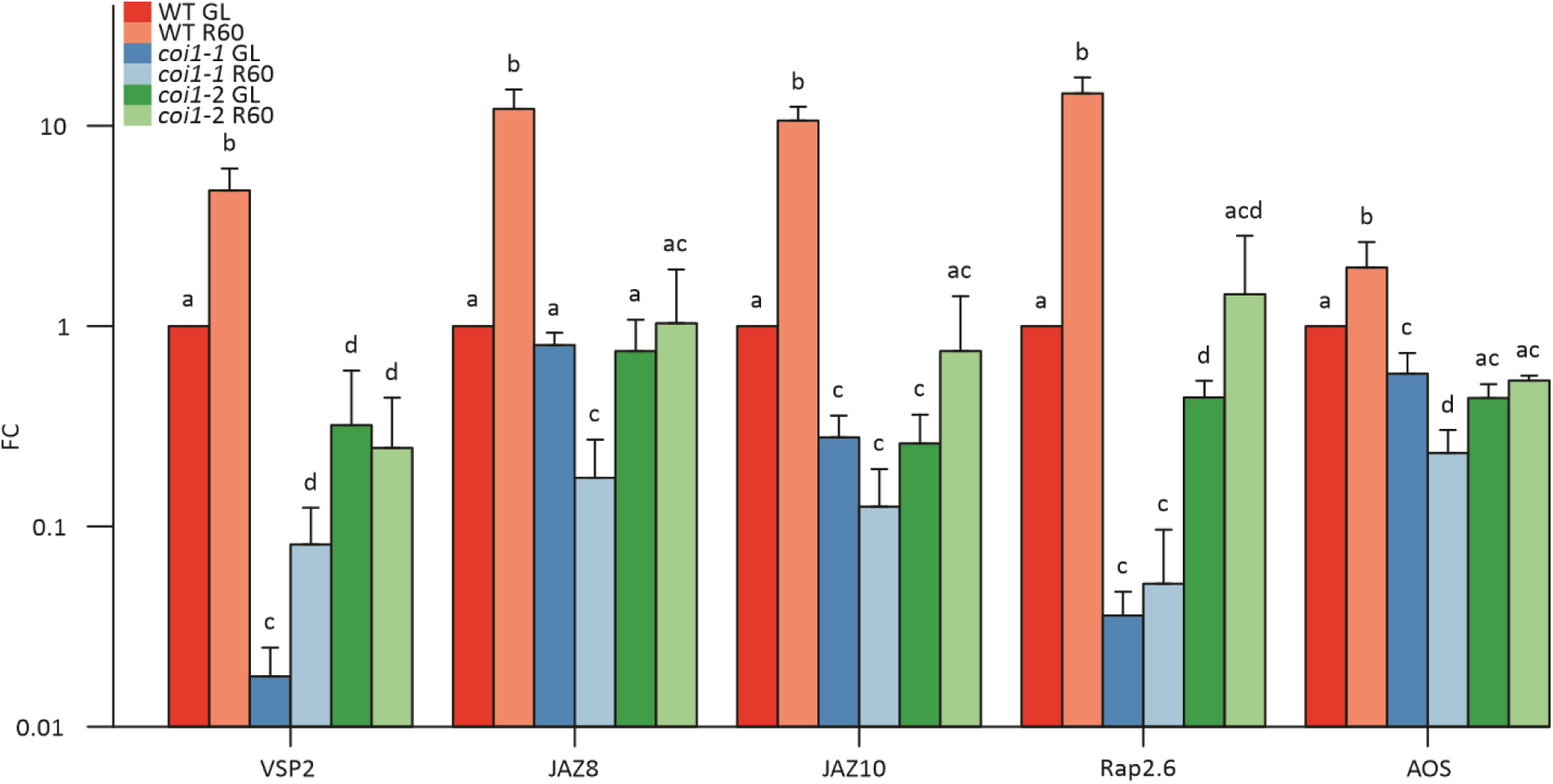
Relative expression of recovery (R)-sustained genes in the wild type (WT) and *coi1* mutants (*coi1-1* and *coi1-2*). qPCR test was performed from leaves illuminated at GL (WT GL; *coi1-1* GL; *coi1-2* GL), and from plants that had been HL-treated for 60 min and subsequently returned to GL for 60 min for recovery from HL stress (WT R60; *coi1-1* R60; *coi1-2* R60). The following R-sustained genes were analyzed: *VSP2*, *JAZ8*, *JAZ10*, *Rap2.6* and *AOS*. The fold change (FC) of gene expression was normalized to WT GL values and expressed as logarithmic scale. Values represent the mean ± SD of three samples. The letters indicate significant differences between the treatments (ANOVA, Tukey-HSD, p < 0.05).

The gene expression profiles in Arabidopsis provided evidence that H_2_O_2_ produced during HL stress may play a role in signaling through induction of the synthesis of OPDA and JA. These oxylipins are central for signaling during the R phase, working in parallel with GSH for balancing the redox state of the cell.

### Short HL exposure is sufficient to enhance the expression of JA/OPDA-responsive recovery genes

We then investigated the duration of the HL pre-treatment required to enhance the expression of R- sustained marker genes in Arabidopsis leaves during the subsequent recovery in GL. For this analysis, the expression of specific R-sustained marker genes (*JAZ8*, *Rap2.6*, *VSP2*, *AOC2*, *AOS*) was assayed with qPCR (Figure 5A). The expression of these genes remained largely unchanged during the HL treatment in comparison to GL, but clear upregulation was detected during the R phase. Even a very short HL pre-treatment of two min was sufficient to induce the upregulation of *JAZ8* and *Rap2.6* genes during the subsequent R phase, whereas *VSP2*, *AOC2* and *AOS* genes required a longer HL pre-treatment of 60 min for upregulation in the R phase (Figure 5A). On the contrary, the expression level of selected *HSP* genes, known to be induced by HL (Table S3), behaved differently. Expression of these genes increased already during the HL but remained stable or decreased during the R phase (Figure 5B).

**Figure 5:**
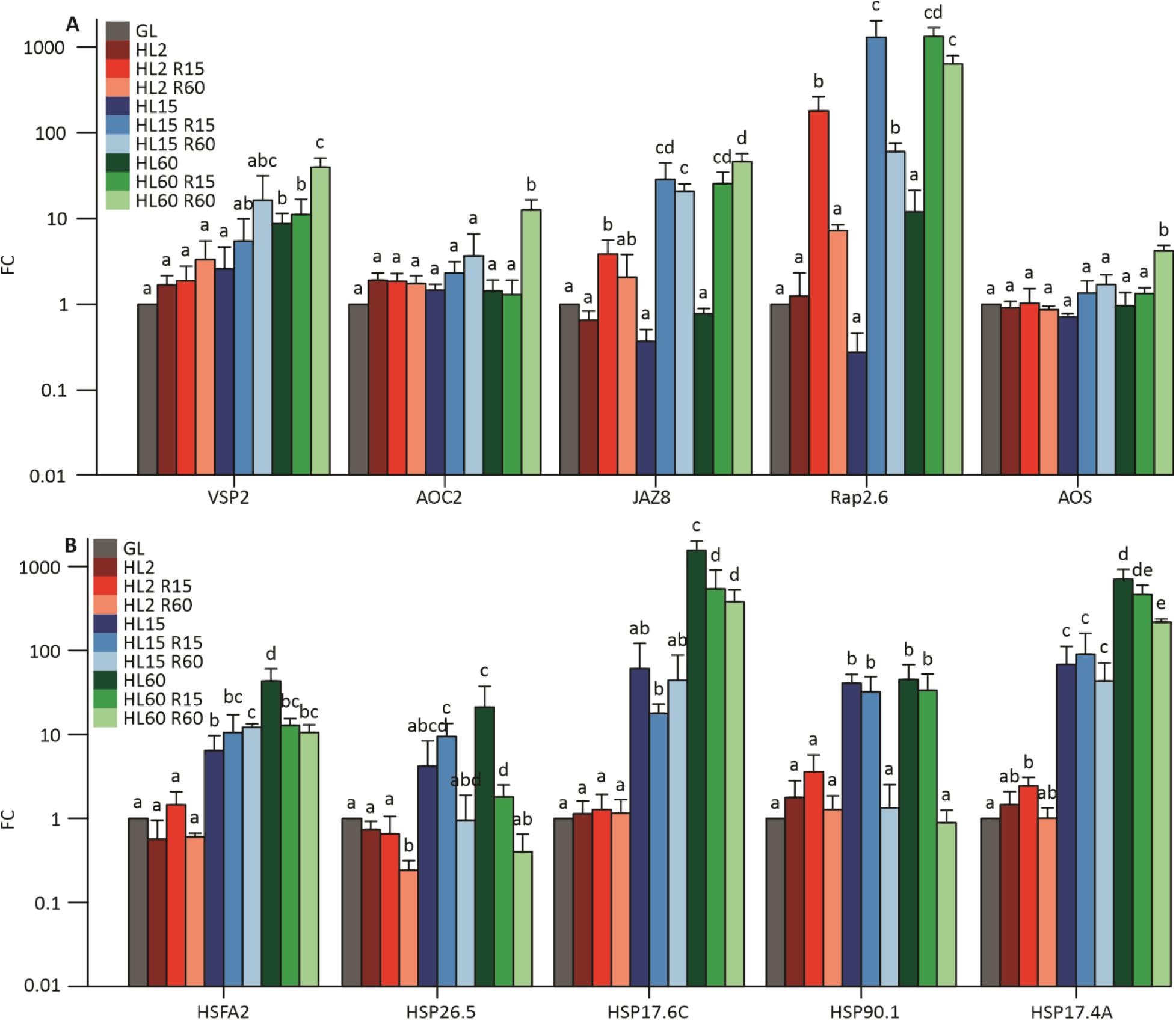
Relative expression of recovery-sustained genes (A) and HSP genes (B) in wild type Arabidopsis leaves exposed to different durations of high light (HL) treatment and subsequent recovery (R) period in growth light (GL). qPCR measurements were done with leaves taken from plants before the HL treatment (GL), and after exposure to HL for 2 min (HL2), 15 min (HL15), or 60 min (HL60). After HL treatment, the leaves were returned to GL and the recovery from HL stress was followed for 15 min (HL2 R15; HL15 R15; HL60 R15) or 60 min (HL2 R60; HL15 R60; HL60 R60). The following R-sustained genes (A): *VSP2*, *AOC2*, *JAZ8*, *Rap2.6* and *AOS*, and HL-induced *HSP* genes (B): *HSFA32*, *HSP26.5*, *HSP17.6C*, *HSP90.1*, and *HSP17.4A*, were analyzed. The fold change (FC) of gene expression was normalized to GL values and expressed as logarithmic scale. Values represent the mean ± SD of four samples. The letters indicate significant differences between the treatments (ANOVA, Tukey-HSD, p < 0.05).

### Changes in leaf hormone and antioxidant contents during HL and recovery phase

To assess the relationship between the gene expression and metabolite accumulation during the HL- exposure and subsequent R phase of the leaves at GL, we next analyzed the abundances of stress hormones, H_2_O_2_, and molecular antioxidants over the course of HL and R. Distinct patterns in metabolite concentrations were displayed throughout the experiment (Figures 6-8 and S4). No significant changes were detected in the amounts of SA and ABA in HL, whereas both ABA and SA decreased during the R phase (Figure 6). OPDA levels displayed an initial rise in HL60, followed by a subsequent decrease during the R phase (Figure 6). In contrast, JA levels clearly increased during the HL treatment, and even more during the subsequent R phase (Figure 6).

**Figure 6:**
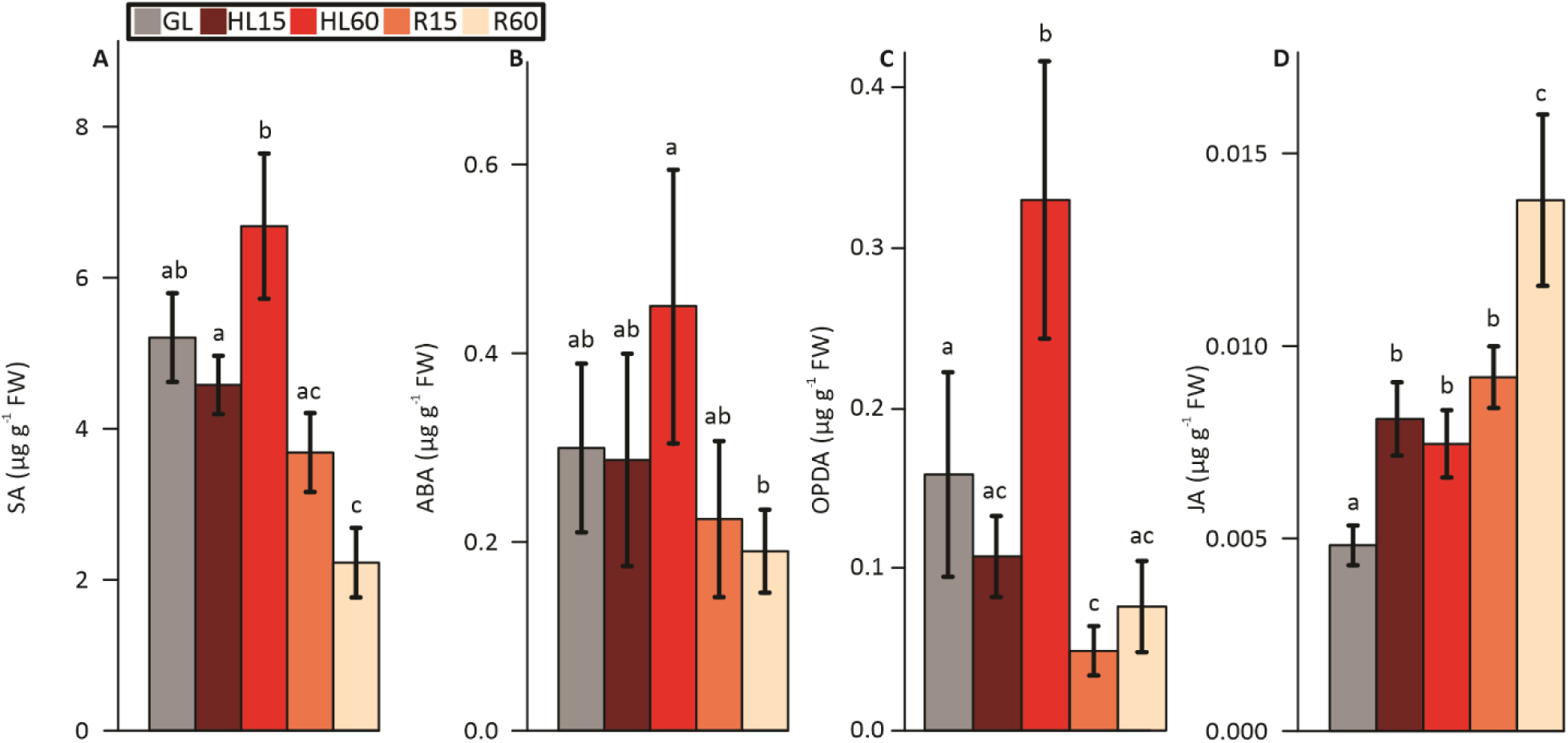
Concentration of A) salicylic acid (SA), B) abscisic acid (ABA), C) 12-oxo-phytodienoic acid (OPDA), and D) jasmonic acid (JA) in leaves exposed to high light (HL) and subsequently transferred to recovery (R) conditions at growth light (GL). Measurements were done with leaves taken from plants before the HL treatment (GL), after 15 min (HL15) and 60 min (HL60) of HL exposure, and during the R at GL for 15 min (R15) and 60 min (R60) after 60 min HL treatment. The values represent the mean ± SD of four samples. The letters indicate significant differences between the treatments (ANOVA, Tukey-HSD, p < 0.05).

The concentrations of molecular antioxidants were also measured to evaluate their roles in response to HL and R treatments. GSH content remained unchanged during the HL treatment but showed a significant increase during the R phase (Figure 7). GSSG peaked in R15 and decreased in R60 (Figure 7). Furthermore, the amount of GS-OPDA conjugate increased during the HL exposure and subsequently decreased during R phase (Figure 7).

**Figure 7:**
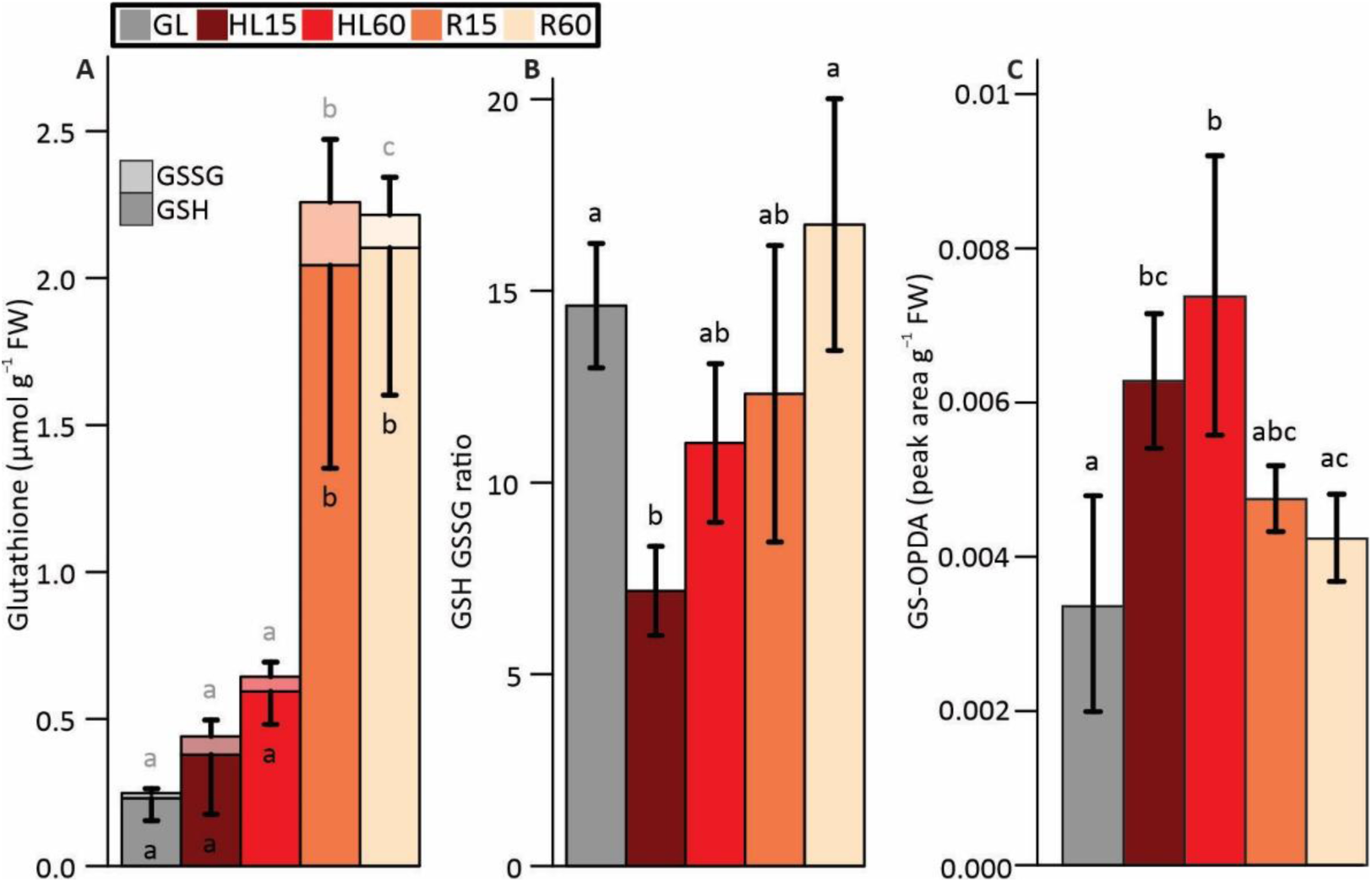
Concentration of glutathione and glutathione-12-oxo-phytodienoic acid (GS-OPDA) conjugates in leaves exposed to high light (HL) and during subsequent recovery (R) at growth light (GL). (A) total glutathione (GSH+GSSG), B) ratio of GSH to GSSG, C) GS-OPDA. Measurements were done with leaves taken from plants before the HL treatment (GL), after 15 min (HL15) and 60 min (HL60) of HL exposure, and during the recovery at GL for 15 min (R15) and 60 min (R60) after the 60 min HL treatment. The values represent the mean ± SD of four samples. The letters indicate significant differences between the treatments (ANOVA, Tukey-HSD, p < 0.05).

Amino acids with antioxidant properties were measured from leaves to assess their contribution to the antioxidant capacity. Methionine levels increased with HL treatment and remained elevated during the R phase (Figure S4). Homocysteine, tryptophan and histidine levels spiked in HL60 and decreased during the R phase (Figure S4).

Reduced ascorbate (AsA) levels increased during the R phase, and accordingly, the levels of oxidized ascorbate (DHA) increased with HL treatment and decreased during R (Figures 8A and B). H_2_O_2_ levels peaked in HL60 and decreased during the R phase, whereas lipid peroxidation levels detected by thiobarbituric acid reactive substances (TBARS), displayed an increase during the R phase (Figures 8C and D).

**Figure 8:**
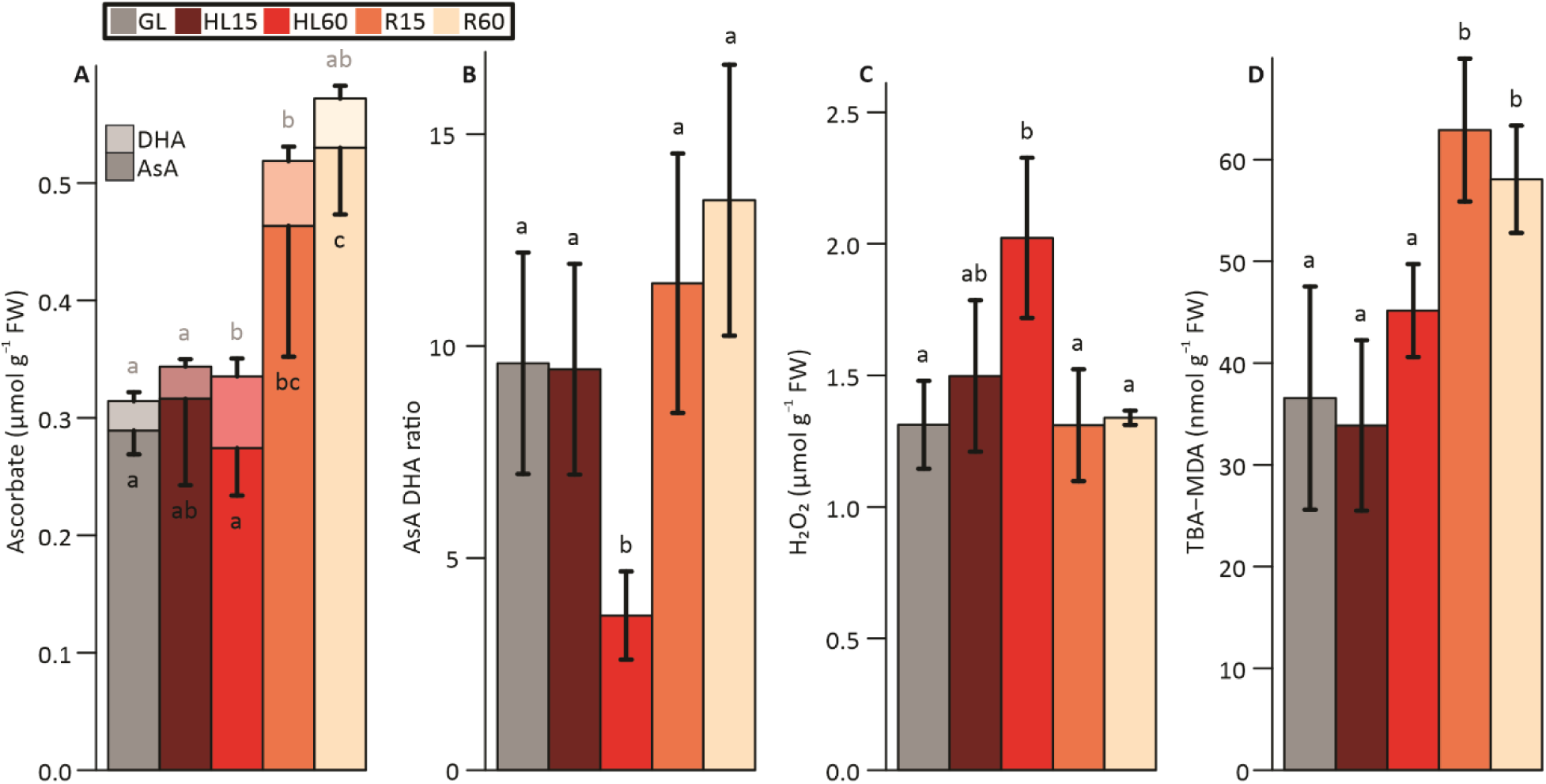
Concentration of ascorbate, hydrogen peroxide (H_2_O_2_), and thiobarbituric acid reactive substances (TBARS) in leaves exposed to high light (HL) and during subsequent recovery (R) at growth light (GL). A) Concentration of total ascorbate (AsA+DHA), B) ratio of AsA to DHA, C) concentration of H_2_O_2_, and D) concentration of TBARS. Measurements were done with leaves taken from plants before the HL treatment (GL), after 15 min (HL15) and 60 min (HL60) of HL exposure, and during the recovery at GL for 15 min (R15) and 60 min (R60) after 60 min HL treatment. The values represent the mean ± SD of four samples. The letters indicate significant differences between the treatments (ANOVA, Tukey-HSD, p < 0.05).

### The *coi1* mutants show slightly faster PSII recovery after the HL treatment

Next, we investigated whether the lack of JA signaling exerts an effect on photosynthetic light reactions in plants exposed to HL and during subsequent recovery at GL. To this end, we made use of the *coi1* mutants that lack JA signaling (Figure 4) and measured the photochemical efficiency of PSII (Fv/Fm) from Arabidopsis leaves. No significant differences in Fv/Fm were detected in HL-exposed plants but during R phase at GL, Fv/Fm recovered slightly faster in the *coi1* mutants (Figure S5).

### HL induces changes in mRNA splicing of the genes

Differential gene expression analysis alone may overlook the complexity of signaling events, especially in the context of alternative gene splicing. Investigating differential isoform usage (DIU) of the genes is crucial as it allows discerning isoform-specific changes, providing a more nuanced understanding of signaling dynamics and cellular responses. DIU analysis revealed 1119 genes exhibiting DIU during the experimental set-up of this article. Most of these genes displayed an increased usage of isoforms that either gained or lost introns (Figure S6). Remarkably, the number of genes gaining introns substantially increased during the HL treatment and the number decreased during the R phase (Figure S6). Furthermore, another set of transcripts had shorter open reading frame (ORF) in HL compared to R phase (Figure S6), which could result in truncated proteins produced in HL-treated leaves.

## Discussion

Relatively short exposures of Arabidopsis rosettes to HL, ranging from minutes to one hour, followed by R periods of 15 or 60 min in GL, were applied here to mimic changes in light intensity under natural conditions. This short HL treatment does not yet induce clear acclimation processes but induces regulatory mechanisms of photosynthesis and metabolic changes that help the plant to survive under frequent changes in light intensity. Photosynthetic regulation mechanisms induced by changes in light intensity, such as state transitions, NPQ, photosynthetic control and cyclic electron flow, have been extensively investigated (Roach and Krieger-Liszkay, 2012; Tikkanen et al., 2015; Yamori et al., 2015), as well as the expression of stress-responsive genes linked to genetic reprogramming in the plant cell during different durations of HL exposure (Gollan et al., 2015; Crisp et al., 2017; Huang et al., 2019; Zandalinas et al., 2020). However, much less is known about the production of signaling metabolites and gene expression profiles related to the R phase after exposure of plants to HL stress (Crisp et al., 2017; Gollan and Aro, 2020). To gain specific information about the R phase, Arabidopsis leaves were subjected to global gene expression analyses and determination of the levels of ROS, stress hormones and antioxidants using the leaves that were (i) kept at GL, (ii) then shifted to HL for short time periods and (iii) finally shifted back to GL for 15 or 60 min for R.

### Short-term HL stress induces hydrogen peroxide, ABA and oxylipin signals with related gene expression changes

To get insights into the triggers underlying gene expression changes during HL stress and R, we first traced the components responsible for initiating the HL-responsive signaling cascades in Arabidopsis leaves in comparison to GL. HL gradually induced H_2_O_2_ accumulation in leaves (Figure 8C), which has generally been reported to be counteracted by production of antioxidants (Gechev et al., 2002; Hernández et al., 2004). Although no significant increase in GSH or ASA contents was evident after HL exposure of Arabidopsis (Figures 7A and 8A), the proportion of oxidized forms of glutathione (GSSG, HL15) and ascorbate (DHA, HL60) increased in leaves, strongly suggesting that GSH and ASA had been utilized for protection against oxidative stress (Figures 7B and 8B), in accordance with earlier reports (Yoshimura et al., 2000; Alvarez-Fernandez et al., 2021). Likewise, an increase in antioxidative amino acids plausibly also contributed to ROS scavenging in HL (Figure S4; Khorobrykh et al., 2020). On the contrary, no signs of lipid peroxidation were evident, detected by TBARS-reactive substances (Figure 8D), thereby eliminating the risk of severe oxidative stress in cells. Instead, it is conceivable that the gradual accumulation of H_2_O_2_ allows its use as a signal to regulate gene expression, inducing protective processes to cut down the further increase in ROS level.

Although the accumulation of H_2_O_2_ was detected only in HL60 leaves (Figure 8C), the upregulation of distinct H_2_O_2_-responsive genes, such as HSPs, was evident after both 15 min and 60 min HL exposure (Figures 3, 5B and S2). These data suggest that H_2_O_2_ is one of the signals produced during leaf exposure to HL, which has been shown to be produced in PSI (Fitzpatrick et al., 2022; Tiwari et al., 2024). Singlet oxygen is also produced in HL (Triantaphylidès et al., 2008; Dmitrieva et al., 2020), while it is very efficiently scavenged by carotenoids and tocopherol (Ramel et al., 2012; Dmitrieva et al., 2020), which is likely reflected in strong differences in ^1^O_2_-responsive gene expression between the 15 and 60 min duration of the HL treatments (Figure S2A). In this work, OPDA and JA biosynthesis genes were moderately or highly upregulated already in HL, and this increased expression was reflected in accumulation of oxylipins in leaves (Figures 1A and 6D), whereas other oxylipin-responsive genes did not substantially change in HL15 but were strongly upregulated in HL60 (Figures 1A and 1B). A portion of ABA-responsive genes was also up-regulated by HL treatment (Figure S3).

Taken together, we conclude that H_2_O_2_, ABA and oxylipins are among the compounds produced by short-term HL treatment and involved in the initiation of respective signaling cascades. This is consistent with previous reports highlighting the role of H_2_O_2_, ABA- and JA- responsive genes in HL stress (Balfagón et al., 2019; Huang et al., 2019; Zandalinas et al., 2020; Alvarez-Fernandez et al., 2021).

### JA signaling gains importance in recovery from short-term HL stress

Considering the signaling pathways in Arabidopsis that mediate the HL- and R-responsive gene expression signals to the nucleus, we focused on the differential accumulation of OPDA and JA (Figure 6) due to their important roles in signaling. OPDA synthesis occurs in chloroplasts and serves as a precursor for JA synthesis, which occurs in peroxisomes (Figure 1; Schaller and Stintzi, 2009; Wasternack and Song, 2017). OPDA accumulation in response to HL treatment has been reported previously (Balfagón et al., 2019), and this was also the case in the current work (Figure 6). However, a strong decrease in OPDA levels is notable in R, strongly suggesting that OPDA was used for JA production, especially in the R phase (Figure 6).

Further analysis of other oxylipin-responsive genes revealed DEGs, whose expression levels specifically paralleled the biosynthesis of OPDA and JA in the leaves during the R phase (Figures 1 and 6). The number of OPDA-induced genes peaked at R15 (46 out of 65 genes; Figure 1C), while the number of JA-induced genes peaked at R60 (57 out of 68 genes; Figure 1B), indicating the stepwise synthesis of JA (Figure 1A). Our results imply that the oxylipin signaling indeed is a crucial component of the R process from HL stress. This conclusion was further strengthened by using the JA receptor *coi1* mutants that showed a lack of expression of R-sustained genes (Figure 4), thereby confirming the involvement JA signaling in R phase. The required duration of the HL pre-treatment was remarkably short since a portion of the R-sustained genes was upregulated in R only after two min of HL pretreatment (Figure 5A). Such a rapid response suggests that even very short exposures to HL, such as canopy movements, may be long enough in duration for accumulation of JA to guide the recovery when the typical canopy conditions have returned.

The primary signal generated by HL that leads to accumulation of JA, both in HL and especially when plants are returned to GL conditions, has remained elusive. H_2_O_2_ has been shown to induce JA biosynthesis (Hieno et al., 2019; Lv et al., 2019), but here we show a decline in H_2_O_2_ levels back to the control level and a decrease in H_2_O_2_ signaling already within 15 min in R conditions (Figure 8C). Furthermore, the expression of the R-sustained genes was not upregulated by H_2_O_2_ treatment per se, but mainly during R after the H_2_O_2_ treatment (Figure 3B), suggesting that H_2_O_2_ is not a primary trigger for R- sustained gene expression but rather induces oxylipin synthesis in treated leaves, which triggers the expression of R-sustained genes. OPDA appears to be a link in accumulation of JA and plant recovery from HL stress. Although the level of H_2_O_2_ decreased during R, lipid peroxidation (accumulation of TBARS), indicating continued ROS production, still continued under R (Figure 8D). The ROS produced at the beginning of R could be ^1^O_2_, since high expression of ^1^O_2_-induced genes was observed at R15 (Figure S2A). Thus, it is possible that OPDA was still produced also during R, but the total amount decreased due to its conversion to JA (Figure 6). Photodamaged PSII complexes could be the source of ^1^O_2_ (Krieger-Liszkay, 2005), since Fv/Fm decreased by about 20 % during HL treatment and damaged PSII complexes were only gradually repaired at GL (Figure S5).

### Glutathione and ascorbate are key antioxidants to guarantee the recovery from HL stress

When Arabidopsis plants were transferred from HL stress to R conditions, a drastic increase in GSH and AsA levels was observed in leaves, and the GSH/GSSG and AsA/DHA ratios were rapidly restored to control levels (Figures 7B and 8B), suggesting a critical role for these antioxidants in R processes. JA has been shown to activate the genes involved in GSH and AsA biosynthesis (Sasaki-Sekimoto et al., 2005), suggesting a synergy between JA signaling and the levels of these antioxidants during recovery. JA has been shown to induce the expression of genes of the sulfur assimilation into Cys, an upstream step in GSH synthesis (Jost et al., 2005; Sasaki-Sekimoto et al., 2005). OPDA is also known to induce GSH production by facilitating CYP20-3 binding to the cysteine synthase complex (CSC) (Park et al., 2013; Liu et al., 2020). The increase in the expression of genes encoding enzymes involved in GSH synthesis (*GSH2*) and recycling (*GGCT2*) correlated with the accumulation of GSH during the R phase (Figure 2). On the other hand, GSH neutralizes the RES activity of OPDA through glutathionylation of unsaturated double bonds (Dorion et al., 2021; Ito and Ohkama-Ohtsu, 2023). Both OPDA and GS-OPDA conjugates accumulated in HL60 and decreased during R phase (Figures 6C and 7C), together with increased *GSH2* and *GGCT2* gene expression and increased GSH levels (Figure 2). The concomitant decreases in OPDA and GS-OPDA conjugate during R suggests that OPDA is both converted to JA and processed in the vacuole by GGT4 (Figure 2).

AsA is another important antioxidant in plant metabolism (Foyer and Noctor, 2011). It is linked to GSH metabolism via the ascorbate-glutathione cycle, which scavenges H_2_O_2_ (Foyer and Noctor, 2011). Like GSH, AsA levels did not increase significantly during the HL exposure, but the content almost doubled in leaves under the R conditions at GL (Figure 8A). In addition, GSH is involved in ROS scavenging via GPXs, PRXs and GRXs (Meyer et al., 2008; Bela et al., 2022). Therefore, it is conceivable that plants do not invest in the synthesis of AsA and GSH during the HL stress and instead favor the production of ROS for signaling purposes. Conversely, when the HL stress is terminated by shifting plants to R conditions, the increase in AsA and GSH synthesis is crucial for the detoxification of ROS (Figure 9).

**Figure 9:**
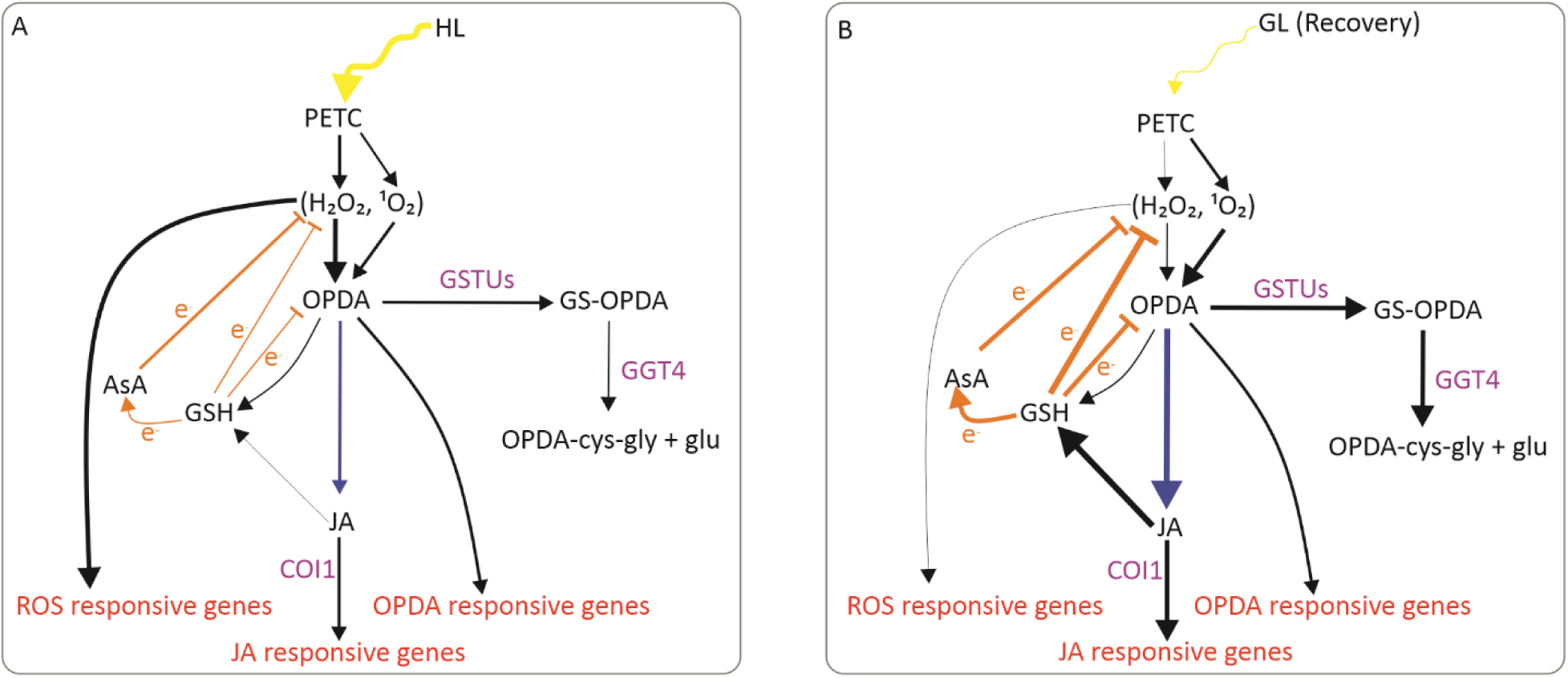
A model of a signaling pathways induced by high light (HL) and during the subsequent recovery at growth light (GL). A) HL triggers the accumulation of reactive oxygen species (H_2_O_2_ and ^1^O_2_), prompting the activation of ROS-responsive genes. Limited levels of reduced glutathione (GSH) and reduced ascorbic acid (ASA) allow ROS signaling to take place. Additionally, ROS activate pathways leading to the synthesis of oxylipins. This results in the accumulation of 12-oxo-phytodienoic acid (OPDA), which is subsequently converted to jasmonic acid (JA). Both OPDA and JA stimulate the expression of oxylipin responsive genes. B) Upon the return to GL for recovery from HL stress, accumulation of H_2_O_2_ diminishes, whereas production of ^1^O_2_ still continues for a while, resulting in continued synthesis of JA via OPDA. OPDA is not accumulating, because in addition to conversion to JA, it is also scavenged by GSH via GLUTATHIONE S- TRANSFERASE (GSTU) activity and the GS-OPDA conjugates are processed in vacuoles by GAMMA- GLUTAMYL TRANSPEPTIDASE 4 (GGT4). OPDA and JA stimulate the synthesis of GSH and ASA, which scavenge ROS and rebalance the redox state of the cells, promoting the recovery. COI1; CORONATINE INSENSITIVE 1. Arrow thickness indicates the intensity of the signaling pathway.

### HL induces intron retention in transcripts

We observed that HL treatment altered RNA splicing of transcripts, which has an additional effect on gene expression. HL treatment led to a significant increase in intron retention in hundreds of transcripts (Figure S6), ultimately altering the translation products. The increase in intron retention in response to light has been reported previously (Zhang et al., 2017; Perrella et al., 2020). It has been suggested that chromatins are more open due to histone acetylation under HL, leading to intron retention as transcription and splicing occur simultaneously (Perrella et al., 2020). The fact that the chromatin is more open means that transcription can occur more quickly, causing some introns to be skipped during splicing.

### Conclusions

Our study provided new insights into the signaling mechanisms used during the R phase after short-term HL stress (Figure 9). H_2_O_2_ accumulates in HL, leading to the synthesis of OPDA, which is used for JA synthesis, especially during the R phase. JA signaling in the R phase induces the accumulation of GSH and AsA, which scavenge the ROS and restore the redox state of the cells, thereby promoting recovery from the stress. Remarkably, even a short exposure to HL stress, as little as two min, is sufficient to initiate some of the signaling cascades typical of the R phase.

## Materials and Methods

### Plant material, growth conditions and light treatments

WT plants of *Arabidopsis thaliana* (L.) Heynh. Columbia (Col-0) ecotype were used to study signaling initiated by HL treatment and in subsequent R phase at GL. WT and *coi1* mutants were used for the study of JA signaling during R from HL stress. *coi1-1* and *coi1-2* mutant lines at Col-0 background were obtained from The Nottingham Arabidopsis Stock Centre (NASC, Nottingham, UK, N68754 and N68755).

Plants were grown under 8h of light (100 μmol photons m^−2^ s^−1^) and 16h darkness for 6 weeks in a phytotron growth chamber. Temperature, CO_2_ concentration and humidity were 23°C, 400 ppm and 60%, respectively. OSRAM PowerStar HQIT 400/D metal halide lamps (Osram, Munich, Germany) were used for illumination during growth. Leaf number seven from Arabidopsis rosettes was used for analysis of metabolites and gene expression.

For transcriptomics and metabolite quantification, WT plants were exposed to 1000 μmol photons m^−2^ s^−1^ for 15 min (HL15) or 60 min (HL60), while subsequent recovery from HL60 involved incubation under GL (100 μmol photons m^−2^ s^−1^) for 15 min (R15) or 60 min (R60).

For qPCR gene expression measurement, leaves of WT plants were harvested from GL and after the HL treatment of 2 (HL2), 5 (HL15) and 60 (HL60) min. Following the HL treatment, plants were transferred to GL for recovery for 15 min (R15) and 60 min (R60).

COI1-dependence of the expression of JA marker genes was assessed with WT, *coi1-1* and *coi1-2* mutant lines. GL samples were harvested before the HL treatment of 60 min, and the R60 samples were harvested from plants transferred to GL for 60 min after the HL treatment.

H_2_O_2_ treatment was done at GL by floating the leaves on 20 mM H_2_O_2_ solution for 5h (H_2_O_2__5h) while leaves of control plants were floated on water for 5h (H_2_O_5h) (Willekens et al., 1997). The recovery from H_2_O_2_ treatment was performed by floating the leaves on water for 5h (H_2_O_2__5h R).

### Measurement of metabolites and glutathione-OPDA conjugate with ultra-performance liquid chromatography

Plant leaves were frozen with liquid nitrogen and ground with mortar to a fine powder. 1 ml of MeOH was added to the leaf powder. The samples were subjected to homogenization for 30 seconds. The suspension was centrifuged at 4°C for 5 minutes at 14000 x g. 800 μl of supernatant was transferred to an Eppendorf tube for ultra-performance liquid chromatography (UPLC) analysis. UPLC analyses were performed by the Turku Metabolomics Centre (Turku Bioscience Centre, Turku, Finland). The standards for hormones and amino acids were obtained from Sigma-Aldrich (Sigma-Aldrich, St. Louis, MO, USA). The concentrations were expressed as μg in g of leaf fresh weight (FW). The standard for glutathione-OPDA (GS-OPDA) conjugate was obtained from the reaction between GSH with OPDA (Dixon and Edwards, 2009) and the concentration was expressed as peak area per g of FW.

### H_2_O_2_ and TBARS measurements

H_2_O_2_ measurement was carried out according to the method described previously (Bela et al., 2018). Leaves were frozen with liquid nitrogen and ground with mortar into a fine powder. 500 μl of 0.1% trichloroacetic acid (TCA) was added to the leaf powder. The suspension was centrifuged at 7000 x g for 20 minutes at 4°C. 250 μl of supernatant was transferred to an Eppendorf tube and 500 μl of 50 mM potassium phosphate buffer (pH 7.0) and 0.5 ml of 1 M potassium iodide were added. Following incubation of 10 min at 25°C, the absorbance of the samples was measured at 390 nm. Concentration of H_2_O_2_ in the sample was calculated by using standard curve with known concentrations of H_2_O_2_ and results expressed as μmol H_2_O_2_ in g of FW.

TBARS were measured as described previously (Lima-Melo et al., 2019) to assess the level of lipid peroxidation as an indicator of oxidative stress. The concentrations were expressed as nmol TBARS in g of FW.

### Glutathione measurement

Glutathione measurements were carried out according to the method described previously (Bela et al., 2018). Plant samples were frozen with liquid nitrogen and ground with a mortar to a fine powder. 1.2 ml of 5% TCA was added to the powder and the suspension was centrifuged at 13,000 x g for 20 min. For the glutathione assay, 100 μl of the supernatant was added to either 100 μl of water (for the total glutathione assay) or 100 μl of 10 % 2-vinylpyridine (for the GSSG assay, to mask GSH). 770 μl of 100 mM phosphate buffer (pH 7.5) containing 10 μl each of 1 mM 5,5’-dithio-bis-(2-nitrobenzoic acid) (DTNB), 1 mM NADPH, and 1 unit of GR (Sigma-Aldrich, St. Louis, MO, USA), was added to the mixture and thoroughly mixed. The absorbance of the reaction mixture was measured at 412 nm. The GSH content was calculated by subtracting GSSG concentration from the concentration of total glutathione. The concentration of glutathione in the sample was calculated by a standard curve that was prepared utilizing solutions of glutathione with known concentrations. The concentrations were expressed as μmol in g of FW.

### Ascorbate measurement

Ascorbate levels were measured as described previously (Zhang et al., 2009). Leaf samples were frozen with liquid nitrogen and ground with mortar to a fine powder to which 1.2 ml of 5% TCA was added. The suspension was centrifuged at 13,000 x g for 20 min. For the determination of total ascorbate, 100 μl of the supernatant were mixed with 100 μl of 10 mM dithiothreitol (DTT). After 10 min incubation, 100 μl of 0.5% N-ethylmaleimide (NEM) was added into the mixture and incubated for 15 min. The suspension was neutralized by adding 50 μl of 1M NaOH followed by an addition of 100 μl of 0.1 M K_3_Fe(CN)_6_ in 50 mM phosphate buffer (pH 7.0) and 1 ml of the 0.1 M FeCL_3_ solution. For reduced ascorbate (AsA), 200 μl of water was added instead of DTT and NEM. The reaction solution was incubated at 37°C for 30 min and the absorbance of the solution was measured at 735 nm. The concentration of oxidized ascorbate (DHA) was calculated by subtracting the concentration of AsA from the concentration of total ascorbate. The concentration of ascorbate in the sample was calculated by a standard curve with known concentrations of ascorbate. The concentrations were expressed as μmol in g of FW.

### RNA isolation and analysis

The leaf samples were frozen in liquid nitrogen and ground in a mortar, followed by RNA extraction using the innuPREP Plant RNA Kit (Analytik Jena, Jena, Germany) according to the kit instructions.

For RNAseq, the RNA extracts were sent to BGI Europe Genomic Center (Copenhagen, Denmark) for sequencing. Transcript reads were quantified with Salmon (v0.12) software using *A. thaliana* genome assembly TAIR10 cDNA sequences for reference. Analysis of differential gene expression were carried out with Bioconductor DESeq2 package (Love et al., 2014). Differentially expressed genes (DEGs) were annotated using the *A. thaliana* genome annotation (TAIR 10.49.gtf). Genes with read counts lower than 10 were eliminated before the differential expression analysis was performed. DEGs with adjusted p-value (padj) < 0.05 were selected to create the Venn diagrams and the heatmaps. Gene enrichment analysis was performed with http://geneontology.org/ software (accessed on 15 June 2023). Genes with expression of −1 ≥ log2(FC) ≥ 1 were selected for gene enrichment and Venn diagram analysis.

For RT-qPCR, iScript™ cDNA Synthesis Kit (Biorad, Hercules, CA, USA) was used to synthesize cDNA from RNA extract. The cDNA solution was diluted fivefold with water before use. RT-qPCR measurements were carried out with a Biorad iq5 real time PCR machine (Biorad, Hercules, CA, USA). RT-qPCR samples contained 10 µl of SYBR Green PCR Master Mix (Biorad, Hercules, CA, USA), 0.5 µl of cDNA, 1 µl of forward primer (10 µM), 1 µl of reverse primer (10 µM) and 3 µl of water. Primers (Table S4) for the reference gene UBIQUITIN CONJUGATING ENZYME 9 (UBC9) and marker genes were designed with QuantPrime (accessed on 10 February 2023) and obtained from Sigma Aldrich (St. Louis, MO, USA).

Isoform switch analysis was performed using a specialized software tool IsoformSwitchAnalyzeR (Vitting-Seerup et al., 2019). IsoformSwitchAnalyzeR uses DEXSeq (Anders et al., 2012) to examine the aligned reads to identify differentially expressed isoforms and conducted statistical analysis to evaluate the significance of isoform switches across various conditions or sample groups. The software identified specific isoforms that exhibited differential usage, along with details on the magnitude and direction of the observed isoform switches. The analyses were used to map alternative splicing events and enrichment of consequences of these events. The quantification of the difference in isoform usage is accomplished by calculating the difference in isoform fraction (dIF), which is computed as the “IF in condition B” - “IF in condition A”.

### Photosynthetic efficiency of PSII

The photosynthetic efficiency of PSII (Fv/Fm) in WT and the *coi1* mutant was measured from the seventh leaf of plants with Dual-PAM-100 (Heinz Walz GmbH, Effeltrich, Germany) according to (Kılıç et al., 2023). The plants were dark acclimated for 20 min before measurements.

## Acknowledgements

We would like to express our gratitude to Ville Käpylä for his assistance with the qPCR experiments.

## Author contributions

M.K., P.J.G., E-M.A., and E.R. designed the research. M.K. and P.J.G performed research. M.K., P.J.G., contributed new analytic/computational/etc. tools; M.K. analyzed data. M.K., P.J.G., E-M.A., and E.R. wrote the paper.

## Supplemental data

The following materials are available in the online version of this article.

**Supplemental Figure S1:** Differentially expressed genes in high light (HL) and during recovery (R) in comparison to growth light (GL).

**Supplemental Figure S2:** Differential expression of singlet oxygen (^1^O_2_) and hydrogen peroxide (H_2_O_2_) - responsive genes in high light (HL) and recovery (R) at growth light (GL) in comparison to GL.

**Supplemental Figure S3:** Differential expression of abscisic acid (ABA) and salicylic acid (SA) responsive genes in high light (HL) and recovery (R) in comparison to growth light (GL).

**Supplemental Figure S4:** Concentration of A) methionine, B) homocysteine, C) tryptophan and D) histidine in leaves exposed to high light (HL) and subsequently moved to recover (R) at growth light (GL).

**Supplemental Figure S5:** Photochemical efficiency of photosystem II (Fv/Fm) in leaves of wild type (WT) and *coi1* mutants (*coi1-1* and *coi1-2*) exposed to the different durations of high light (HL) and under subsequent recovery (R) periods at growth light (GL).

**Supplemental Figure S6:** Alternative splicing events in high light (HL) and during the recovery (R) from HL in comparison to growth light (GL).

**Supplemental Table S1:** Gene enrichment analysis of biological processes upregulated in high light (HL) and recovery (R) at growth light (GL) in comparison to GL control conditions.

**Supplemental Table S2:** List of the genes presented in Figure 2.

**Supplemental Table S3:** Average transcript counts of marker genes in RNAseq analyses selected for the qPCR test presented in Figure 3 and Figure 4.

**Supplemental Table S4:** Marker genes and the primers used in the RT-qPCR test.

## Data availability

The data supporting the findings of this study are available in the article and its supplementary materials. The transcriptome data have been deposited to the NCBI GEO repository under GSE277977.

